# FL-QSAR: a federated learning based QSAR prototype for collaborative drug discovery

**DOI:** 10.1101/2020.02.27.950592

**Authors:** Shaoqi Chen, Dongyu Xue, Guohui Chuai, Qiang Yang, Qi Liu

## Abstract

**Motivation:** Quantitative structure-activity relationship (QSAR) analysis is commonly used in drug discovery. Collaborations among pharmaceutical institutions can lead to a better performance in QSAR prediction, however, intellectual property and related financial interests remain substantially hindering inter-institutional collaborations in QSAR modeling for drug discovery.

**Results:** For the first time, we verified the feasibility of applying the horizontal federated learning (HFL), which is a recently developed collaborative and privacy-preserving learning framework to perform QSAR analysis. A prototype platform of federated-learning-based QSAR modeling for collaborative drug discovery, i.e, FL-QSAR, is presented accordingly. We first compared the HFL framework with a classic privacy-preserving computation framework, i.e., secure multiparty computation (MPC) to indicate its difference from various perspective. Then we compared FL-QSAR with the public collaboration in terms of QSAR modeling. Our extensive experiments demonstrated that (1) collaboration by FL-QSAR outperforms a single client using only its private data, and (2) collaboration by FL-QSAR achieves almost the same performance as that of collaboration via cleartext learning algorithms using all shared information. Taking together, our results indicate that FL-QSAR under the HFL framework provides an efficient solution to break the barriers between pharmaceutical institutions in QSAR modeling, therefore promote the development of collaborative and privacy-preserving drug discovery with extendable ability to other privacy-related biomedical areas.

**Availability and implementation:** The source codes of the federated learning simulation and FL-QSAR are available on the GitHub: https://github.com/bm2-lab/FL-QSAR

## 1 Introduction

During the drug discovery process, predicting and prioritizing the properties of large numbers of compounds is an essential step in the early stage of drug discovery (Vamathevan, et al., 2019). In the pharmaceutical industry, QSAR is a commonly used in-silico technique to predict and investigate various properties of compounds, such as the compound affinity towards a target and the absorption, distribution, metabolism and excretion (ADME), etc, which can substantially reduce the experimental work needed here (Ma, et al., 2015).

Machine learning (ML) approaches provide powerful tools that can promote the data-driven decision making, speed up the process and reduce failure rates with abundant, high-quality data in drug discovery and development (Cohen, et al., 2018; Cruz-Roa, et al., 2017; Ding, et al., 2018; Rahman, et al., 2017; Wang and Gu, 2018). Generally speaking, increasing the training data can improve the performance of ML approaches. In the pharmaceutical industry, generating more data generally means more time and money costs (Ma, et al., 2020), however, the structure of the lead compounds in the development pipelines will not be exposed before marketing, which prevents the compound data sharing from one institute to each other. Therefore, although collaboration and data sharing between individual entities is expected to serve as a good strategy to save cost and promote drug discovery, such forms of collaboration have been limited by concerns on compound intellectual property and other related financial interests (Hie, et al., 2018).

Modern cryptography has provided partial solutions to this issue. Secure multiparty computation (MPC) is an increasing popular while classical encryption method that allow multiple entities to compute over their private datasets without revealing any information about pharmacological data (Aho, 1987; Yao, 1982). A quick while thorough survey indicates that MPC has been applied for genomic diagnosis (Jagadeesh, et al., 2017), Drug-target interaction (DTI) prediction (Hie, et al., 2018; Ma, et al., 2020) and genome-wide association study (GWAS) (Cho, et al., 2018). In the MPC framework, all participants must submit their data securely to the third party with encryptions (Bogdanov, et al., 2008). However, participants may not want to share data with the third party, even though the data is encrypted. Meanwhile, other ethic and political issues remain under the MPC framework since countries around the world are strengthening laws to protect data privacy and security by prohibition of certain data transition across countries or organizations, even though the data is encrypted. Such regulations includes the GDPR implemented by the European Union (Voigt and Von dem Bussche, 2017) and CCPA enacted by California, U.S. (de la Torre, 2018) et c. Therefore, the traditional MPC framework remains facing the challenges under the constantly emerging new data laws and regulations.

Federated learning (FL) is a recently proposed collaborative paradigm to enable the data owners collaboratively train a model while any data owner does not expose its data to others (Kairouz, et al., 2019). FL was first proposed by Google (Konečný, et al., 2016; Konečný, et al., 2016; McMahan, et al., 2016), and then extended by Yang et al (Yang, et al., 2019). FL can be categorized into horizontal federated learning (HFL), vertical federated learning(VFL) and federated transfer learning(FTL) (Yang, et al., 2019). HFL is applicable to the scenarios that data sets share the same feature space but differ in samples. VFL applies to the scenarios that data sets share the same sample space but differ in feature space. FTL is introduced in the scenarios that the two data sets differ not only in samples but also in feature space with only a small portion of the feature space and sample space overlapped (Yang, et al., 2019) (**Fig. 1**). In the pharmaceutical industry, the most common scenario is that different institutions often have the same type of data, i.e., the lead compounds in their pipelines, which can be encoded with the same feature representations. The difference is that the compounds held by different institutions are different. Compared to VFL and FTL, HFL is obviously more suitable for the collaborations among pharmaceutical industry. Essentially, HFL is based on the traditional MPC framework with advantages that it passes encrypted model parameters to the server instead of encrypted raw pharmacological data, which provides a workable solution to the facing issues aforementioned.

**Fig. 1.**
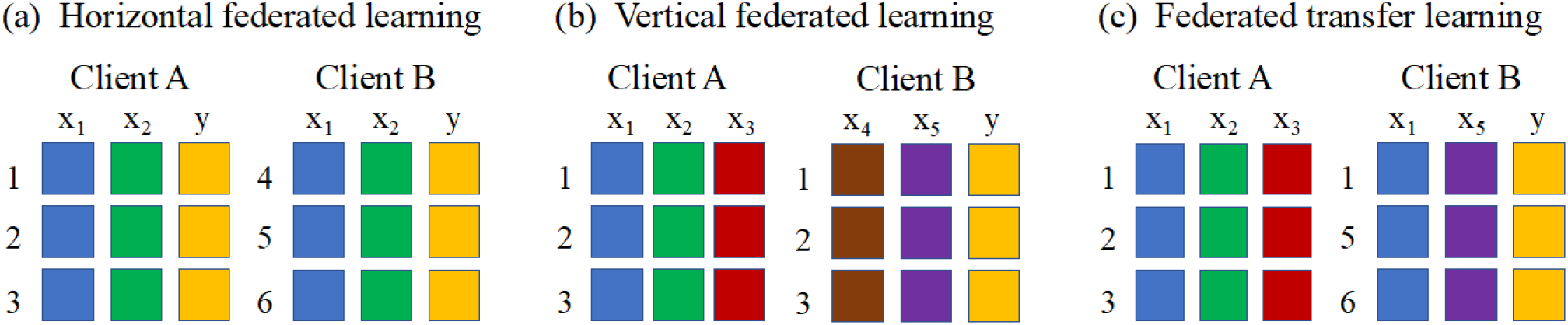
Categorization of Federated Learning. (a) Horizontal federated learning. (b) Vertical federated learning. (c) Federated transfer learning. The term *x_n_* donates the *n_th_* feature of the feature descriptions of the sample, *y* is the sample label and *n* donates sample ID.

In this study, we verified the feasibility of applying HFL to collaborative drug discovery, with the commonly faced QSAR modeling as a demonstration study. To the best of our knowledge, this is the first study to apply FL framework for collaborative drug discovery compared to traditional MPC ones. In our study, we simulated the scenario that parties have their private data, respectively, and trained a routine ML model with and without HFL. Comprehensive experiments indicated that in most cases, collaboration via HFL achieves almost the same performance as that of collaboration via the corresponding cleartext algorithm and significantly outperforms a single client via the corresponding cleartext algorithm. The FL-QSAR platform, which is designed as a prototype system for QSAR modeling under HFL is presented, served as a pioneer study to call for more attention and devoting in this area. To sum up, for the first time, our study demonstrated the effectiveness of applying HFL in QSAR modeling and proposed a prototype framework FL-QSAR with extendable ability. The FL framework and FL-QSAR developed in our study can be applied or extended to various drug-related learning problems involving collaboration and privacy-preserving, promoting the development of collaborative drug discovery and privacy-related computing in pharmaceutical community.

## 2 Methods

### 2.1 Traditional Secure MPC

Secure MPC is a designed protocol that allows multiple parties to compute a function on encrypted data and access only their own data and data that all parties agree to reveal. In secure MPC, the data owner encrypts its data by splitting it using random masks into *n* random shares that can be combined to reconstruct the original data. These *n* shares are then distributed between *n* parties. This process is called secret sharing (Ben-Or and Wigderson, 1988). We use a concrete example to better illustrate the secret sharing concept. Suppose that an integer *x* is the private data and *Q* is a big random integer. Then this client sends a random integer *a* between −*Q* and *Q* to party *S*_1_ and (*x* − *a* mod *Q*) to party *S*_2_ respectively, *a* and (*x* − *a* mod *Q*) are called the two shares of *x* respectively. Then the parties can compute functions on the data by operating on the secret shares and they can decrypt the final result by communicating the resulting shares among each other (Ma, et al., 2020).

### 2.2 Horizonal federated learning applied in FL-QSAR

We then introduced the HFL framework applied in FL-QSAR, as an alternative strategy for collaborative and privacy-preserving QSAR modeling compared to MPC. The formal description of HFL is given as follows:

Define *N* data owners {*F*_1_,…*F_N_*}. All of them would like to train a model by consolidating their respective data {*D_1_*,…*D_N_*}. A conventional method, such as using the traditional MPC, is to put their data together and use *D* = *D*_1_ ∪ … ∪ *D_N_* to train a model. A FL system is a process in which the data owners collaboratively train a model and no data owner *F_i_* expose its data *D_i_* to others. HFL is applicable when data sets share the same feature space while different samples. We denote the features space of the sample as *X*, the label space of the sample as *Y* and the sample ID space as *I*, all of whom constitute the complete training dataset (*I*, *X*, *Y*). HFL can be represented as:

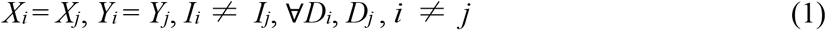

A HFL system commonly assumes *honest* participants and security against an *honest-but-curious* server. An honest client means the client will not send fake or false data to the server. An honest-but-curious server means the server will not send fake or false data to clients, but is curious about information of clients and will mine sensitive information from the data (Yang, et al., 2019).

As shown in **Fig. 2**, in HFL system, participants with the same data structure learn a ML model collaboratively with the help of a server. To train a model to predict QSAR with HFL, our training process of such a system can be divided into the following four steps (Yang, et al., 2019):

- Step 1: Each participant trains its model on its own data, encrypts model parameters with secure MPC techniques, and sends encrypted results to server;
- Step 2: Server performs secure aggregation without obtaining information from any participant;
- Step 3: Server sends back the aggregated results to participants;
- Step 4: Participants update their models with the decrypted parameters.

**Fig. 2.**
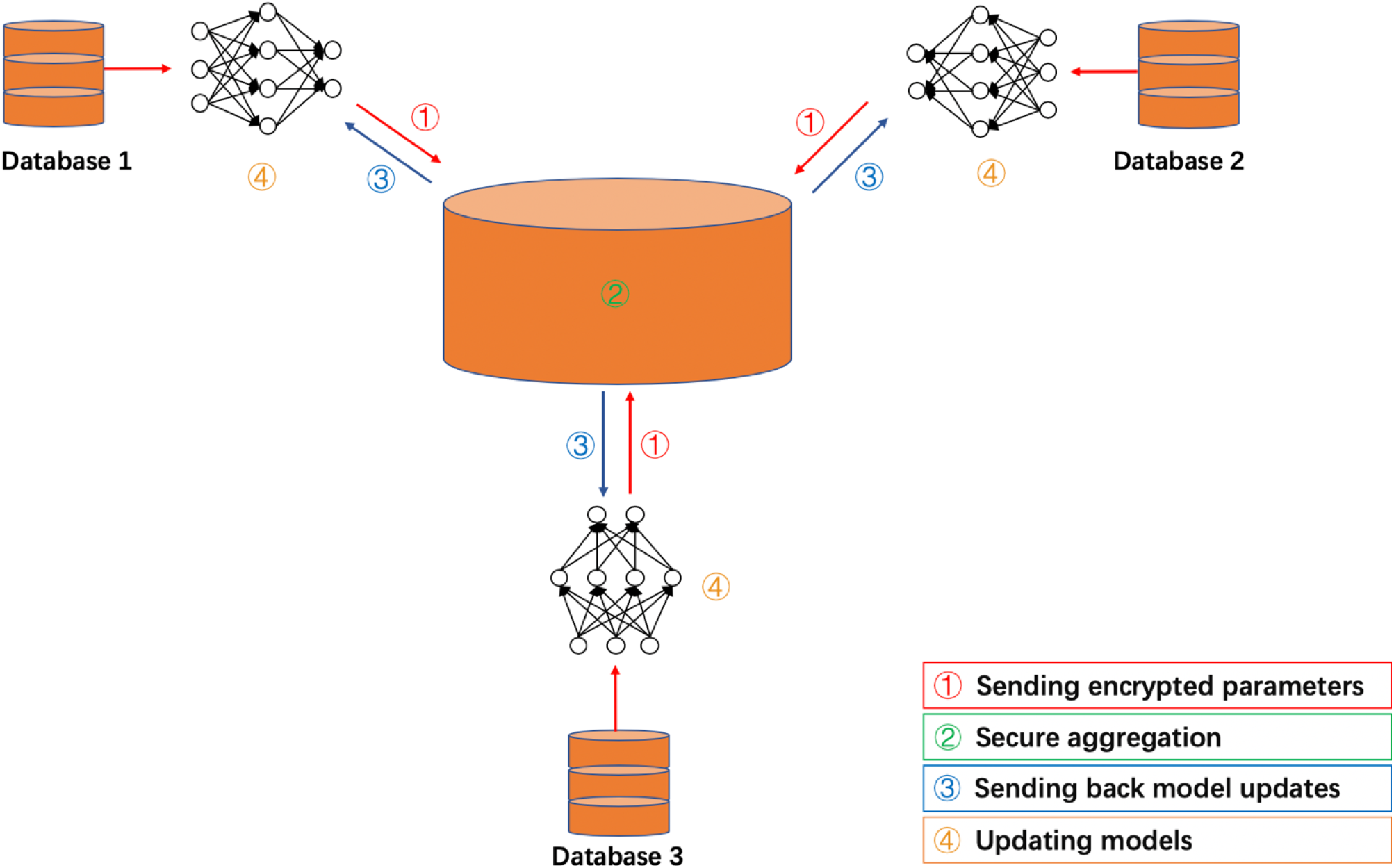
The workflow of HFL. Each client trains its model on its private data, encrypts model parameter, and sends them to the server. The server performs secure aggregation and sends back the aggregated results to clients. At last, clients update their models with the decrypted parameters.

In our implementation of FL-QSAR, we use the Crypten, which is a ML framework built on PyTorch (Paszke, et al., 2019) by applying secure MPC as its cryptographic part, to implement encryption of model parameters and their operations in secure aggregation.

It should be noted that traditional MPC allows the encrypted parameters to compute in a secure manner, and a solution is needed to aggregate these parameters to form the parameters in a federated learning model. One of the most common solutions to aggregation for FL is the Federated Averaging algorithm (McMahan, et al., 2016) (Algorithm *S*1). In our study, the basic idea of the algorithm is to average the parameters *w* and *b* of the neural network models applied in FL-QSAR for aggregating different clients.

**Algorithm S1.**
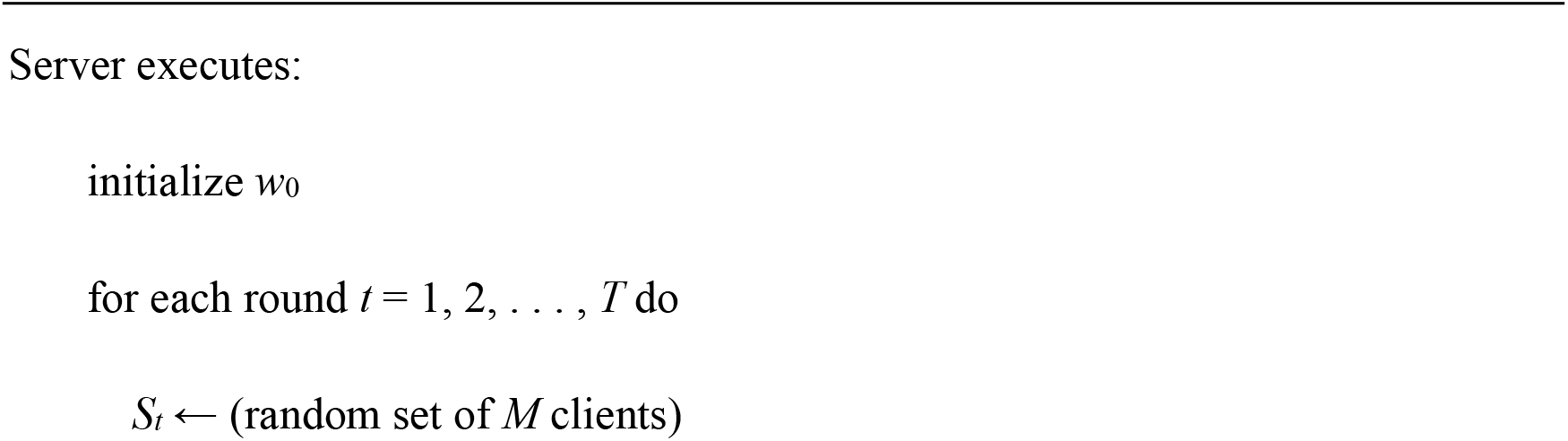
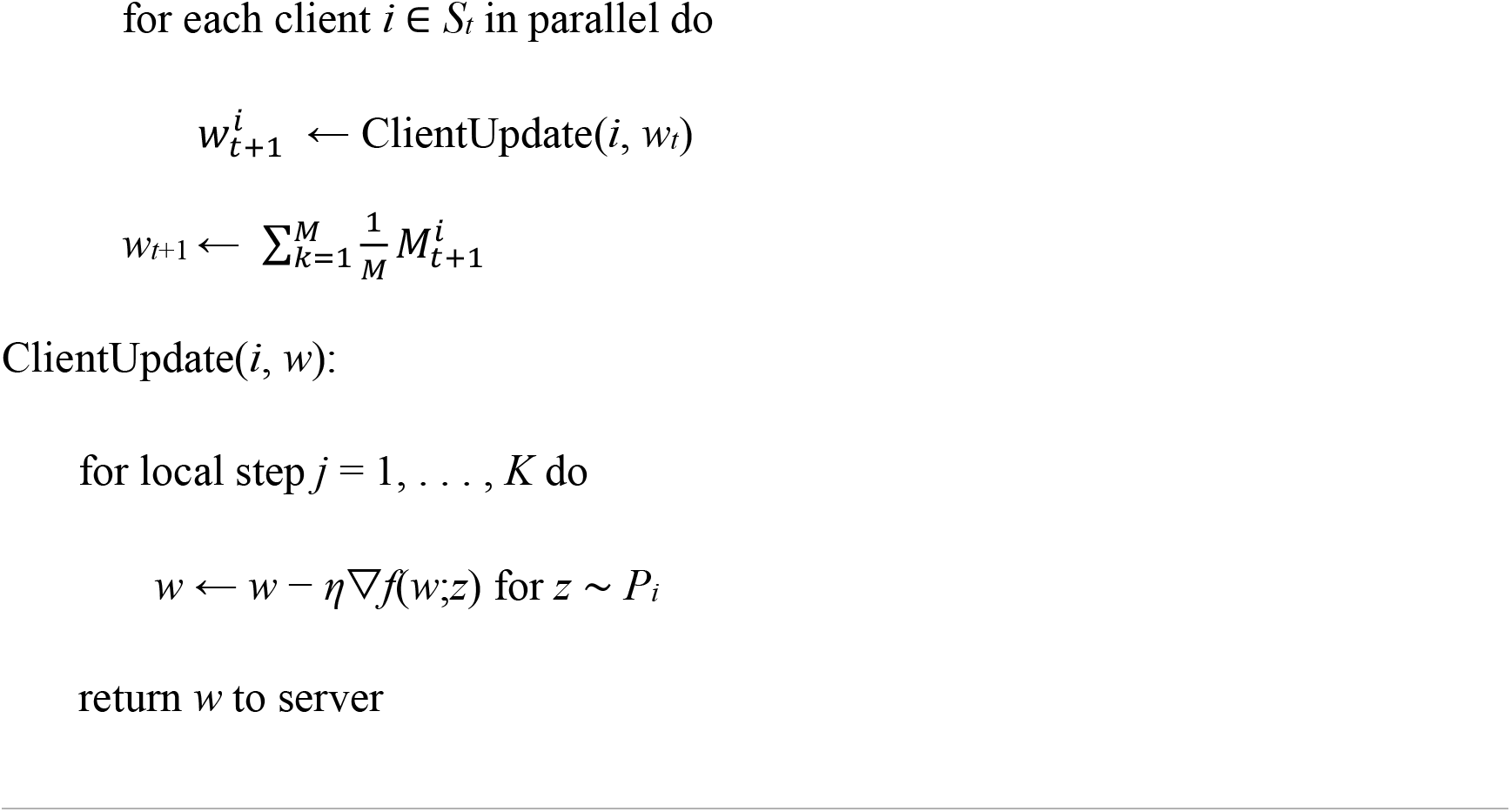
Federated Averaging. The *K* clients are indexed by *k*, *T* is total communication rounds, *M* is the number of local epochs, and *η* is the learning rate.

### 2.3 Benchmark data collection and federated learning simulation for FL-QSAR

In our study, we simulated the scenarios of two clients, three clients and four clients with and without HFL, respectively, to demonstrate the effectiveness of FL-QSAR in QSAR modeling. We take the simulation for 3 clients with and without HFL as an illustration example to demonstrate the process of simulation (**Fig. 3**). We use 15 benchmark datasets public available in Kaggle competition (Ma, et al., 2015) to validate our study. Each dataset is curated for a type of ADME assays or a target, and the detailed description of the datasets are listed in Table 1. Since the performance of the HIVPROT dataset in 15 datasets is extremely outrageous and unstable, it is excluded in the subsequent analysis. We randomly separated all available data into subsets and distributed them to clients, which were then regarded as the private data. Clients have the same amount of training data and testing data, with the chemical structure descriptors (Ma, et al., 2015) as feature descriptions and bioactivities as the labels for the training data. Suppose that there are *n* instances in all available training data. Then a client owns a random subset of training data with 1/*x* · *n* instances, where *x* stands for the number of clients.

**Table 1.**
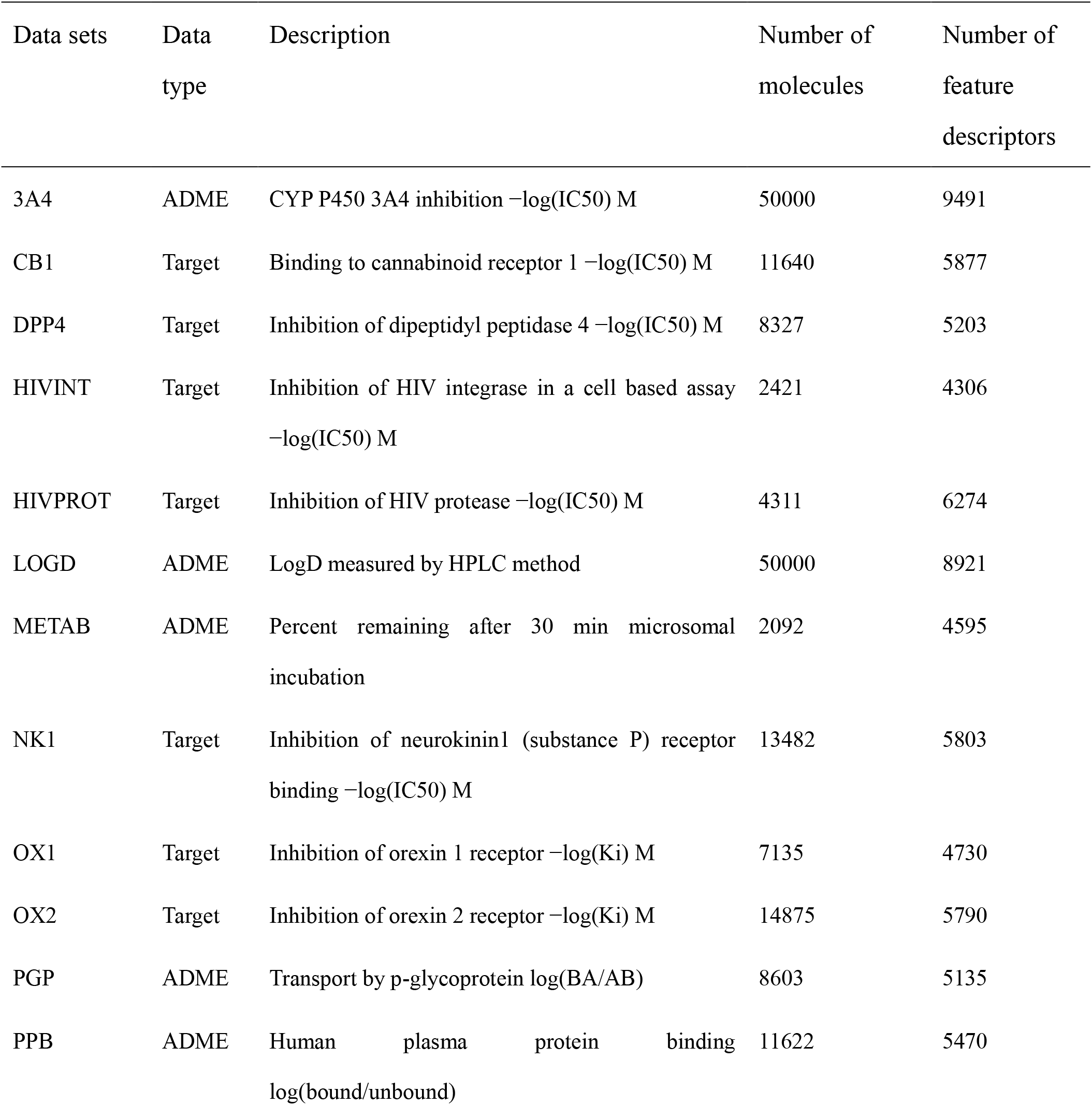
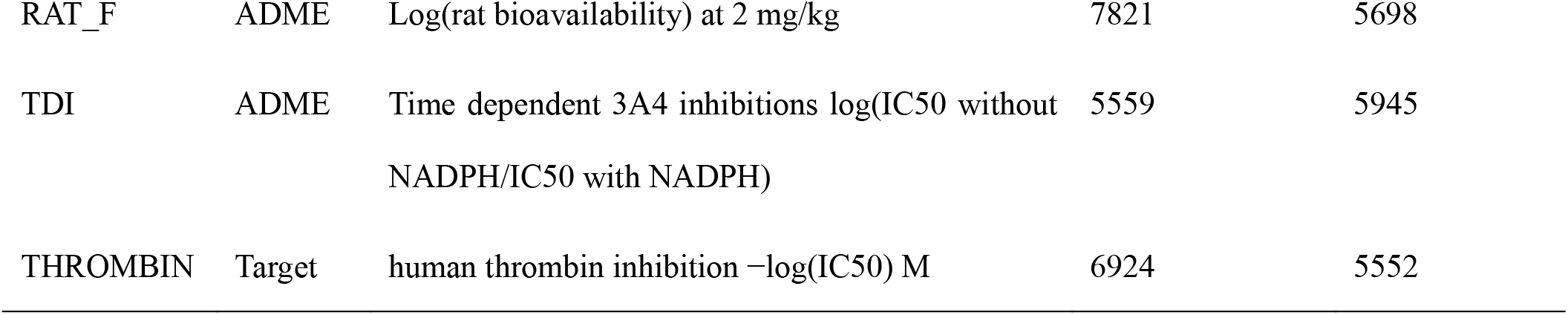
Benchmark data sets tested in FL-QSAR.

| Data sets | Data type | Description | Number of molecules | Number of feature descriptors |
| --- | --- | --- | --- | --- |
| 3A4 | ADME | CYP P450 3A4 inhibition $-\log(\text{IC}_{50})$ M | 50000 | 9491 |
| CB1 | Target | Binding to cannabinoid receptor 1 $-\log(\text{IC}_{50})$ M | 11640 | 5877 |
| DPP4 | Target | Inhibition of dipeptidyl peptidase 4 $-\log(\text{IC}_{50})$ M | 8327 | 5203 |
| HIVINT | Target | Inhibition of HIV integrase in a cell based assay $-\log(\text{IC}_{50})$ M | 2421 | 4306 |
| HIVPROT | Target | Inhibition of HIV protease $-\log(\text{IC}_{50})$ M | 4311 | 6274 |
| LOGD | ADME | LogD measured by HPLC method | 50000 | 8921 |
| METAB | ADME | Percent remaining after 30 min microsomal incubation | 2092 | 4595 |
| NK1 | Target | Inhibition of neurokinin1 (substance P) receptor binding $-\log(\text{IC}_{50})$ M | 13482 | 5803 |
| OX1 | Target | Inhibition of orexin 1 receptor $-\log(\text{K}_i)$ M | 7135 | 4730 |
| OX2 | Target | Inhibition of orexin 2 receptor $-\log(\text{K}_i)$ M | 14875 | 5790 |
| PGP | ADME | Transport by p-glycoprotein $\log(\text{BA}/\text{AB})$ | 8603 | 5135 |
| PPB | ADME | Human plasma protein binding $\log(\text{bound}/\text{unbound})$ | 11622 | 5470 |
| RAT_F | ADME | Log(rat bioavailability) at 2 mg/kg | 7821 | 5698 |
| TDI | ADME | Time dependent 3A4 inhibitions log(IC50 without NADPH/IC50 with NADPH) | 5559 | 5945 |
| THROMBIN | Target | human thrombin inhibition $-\log(\text{IC}_{50})$ M | 6924 | 5552 |

**Fig. 3.**
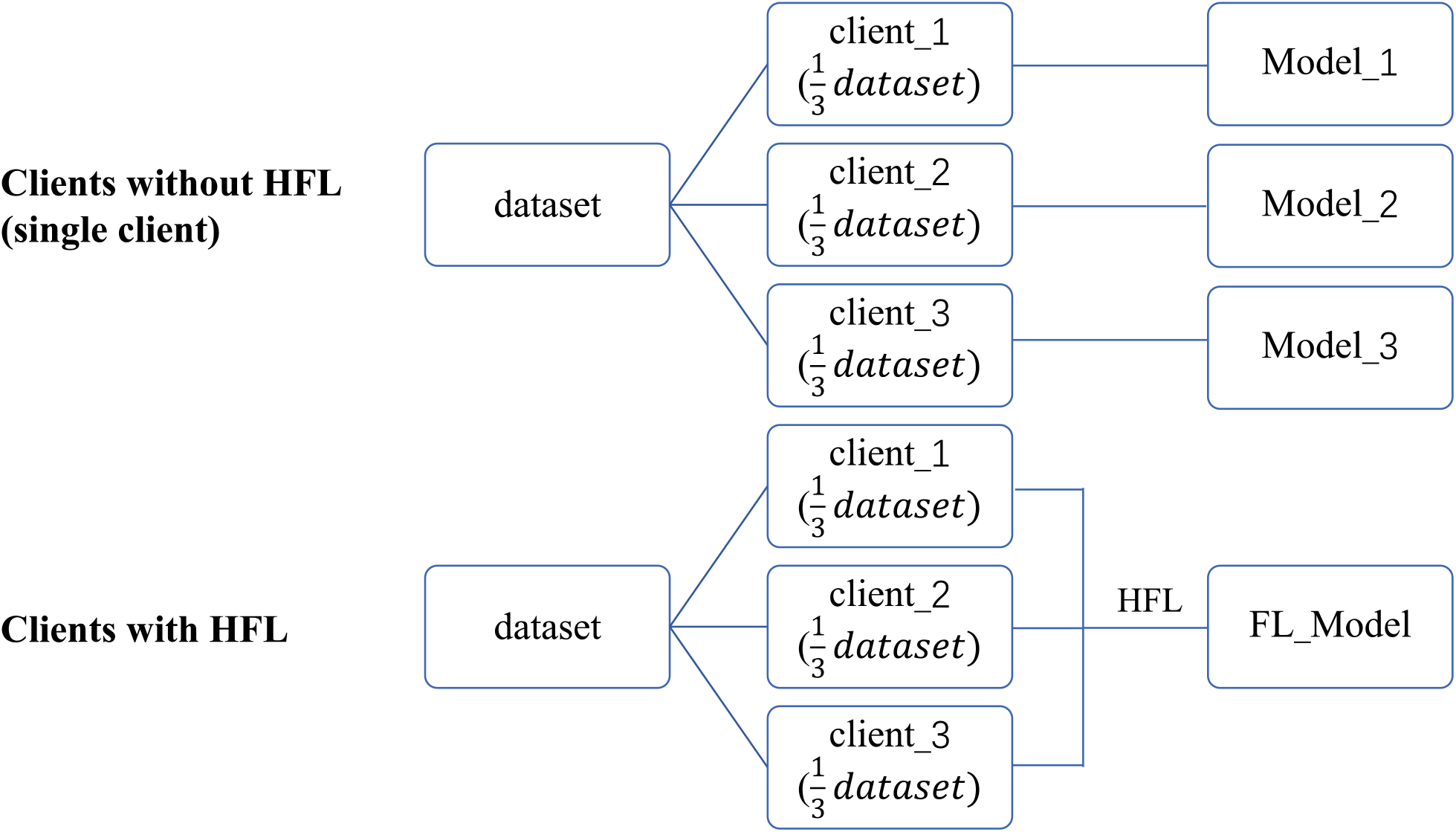
The simulation for 3 clients with and without HFL as a demonstration example

Since the deep neural network has been extensively applied in QSAR modeling and proven to outperform other traditional ML methods (Preuer, et al., 2018), in our study, we just introduced a routine neural network for QSAR model and use parameter setting of DNN recommended by previous study as our hyperparameters in all experiments (Ma, et al., 2015). We consider chemical structure descriptors as the feature descriptions here and the numerical bioactivities are taken as the labels in FL-QSAR, therefore the whole QSAR modeling is taken as a regression problem. In order to improve the numeric stability, logarithmic transformation is applied to transform the features while Min-Max Normalization is performed to scale the labels. The number of training epochs is set to 180 at most and early stopping is adopted to prevent overfitting. In our experiments, we trained the neural networks ten times and averaged their predicted scores as the final results. It should be noted that basically the core ML model applied in FL-QSAR and model selection procedure are not our focus here, and users can try other ML models with different parameter settings and different compound feature descriptors in the future.

## 3 Result

### 3.1 Comparisons between HFL and MPC

We made a comprehensive comparison between HFL and MPC, as MPC was also previously proposed for QSAR modeling (Ma, et al., 2020) (**Table 2)**. First, HFL is a framework developed based on MPC and MPC is used to implement the encryption of model parameters as well as their operations in secure aggregation in the workflow of HFL. Second, MPC combines encrypted raw data of all clients together to train a model while HFL aggregates encrypted model parameters for all clients to train a model. Third, in terms of prediction performance, MPC is just identical to the corresponding cleartext neural network since it essentially trains the model by combining all the training data from individual client, while HFL is expected to be a little bit lower, keep a tradeoff between model efficiency and data private-preserving. Therefore, the upper-bound of the performance of FL-QSAR is the performance of cleartext QSAR modeling by combining all the training data from individual client. Taking together, HFL can solve the issues or concerns by facing the increasing emerged data laws and regulations to prohibit data transmission, where MPC is not able to, and at the same time, reach relatively the same performance as those of combining individual client together for model training.

**Table 2.**
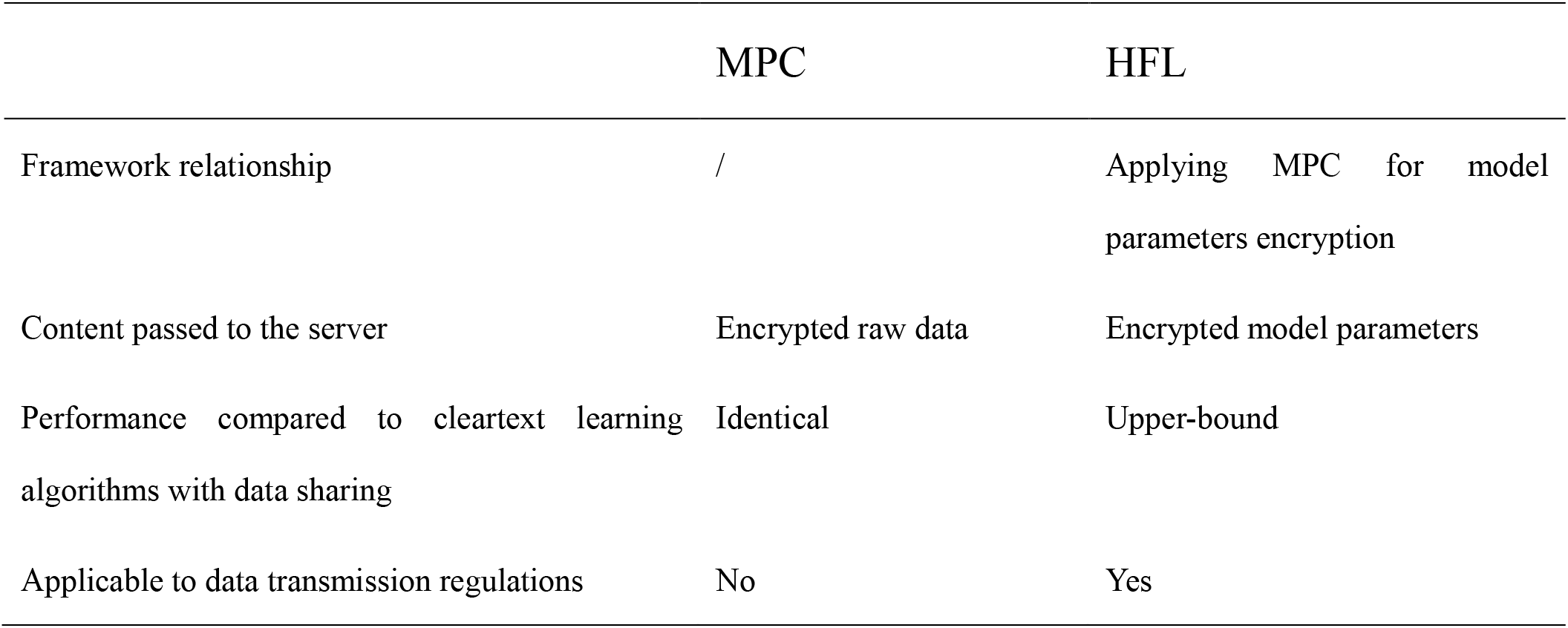
A comprehensive comparison between MPC and HFL.

### 3.2 Comparisons between public and HFL collaborations

In this section, we examined whether collaboration via HFL will cause the loss of the prediction accuracy and to what degree it will cause when compared to the collaboration via cleartext learning algorithms using all shared information. We used the coefficient of determination (R^2^) as the criterion to evaluate the performance for this regression task of QSAR modeling. As shown in Fig. 4a, it indicated that the loss of the prediction accuracy is increasing as the number of clients increases, and HFL for 4 clients is more unstable as 2 outliers appeared. This is expected, since increasing the number of clients will not increase the whole training data, while it increases the model instability because the errors occurred in every single client will influence the final aggregation results in QSAR modeling. Nevertheless, in most cases, HFL can achieve almost the same performance as collaboration via cleartext learning algorithms using all shared information. This can be seen in Fig. 4a that the loss for all scenarios are nearly zero. More details can be found in the Supplementary Table 1.

**Fig. 4.**
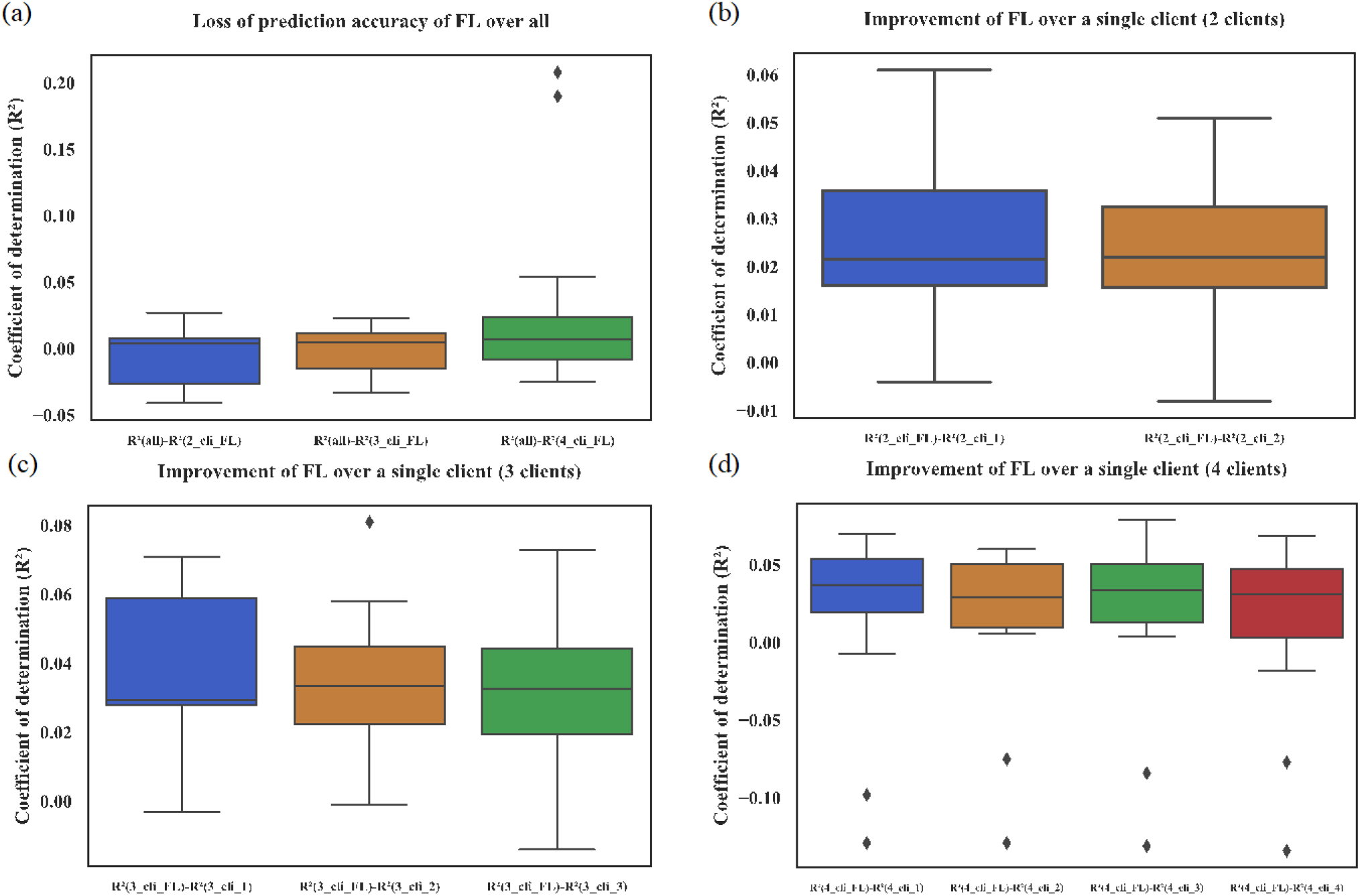
The prediction performance of HFL. The term ‘all’ means that collaboration via cleartext learning algorithms using all shared information. The term ‘*n*_cli_FL’ means that collaboration of *n* clients via HFL and so on. The term ‘*n*_cli_*p*’ means that the *p_th_* client of *n* clients and so on. (a) The loss of prediction accuracy of FL over all. (b) The improvement of HFL for 2 clients over a single client. (c) The improvement of HFL for 3 clients over a single client. (d) The improvement of HFL for 4 clients over a single client.

### 3.3 Comparison between HFL collaborations and a single client using private data

We also compared the prediction performance between a single client and that of HFL for 2 clients, 3 clients and 4 clients respectively. We found that collaboration by HFL achieved a significant improvement in R^2^ and outperformed a single client using its private data (**Fig. 4b-4d**). This result demonstrated the effectiveness of HFL. Collaboration by HFL gained a substantially performance improvement than that of one client by using only its private data. More details can be found in the Supplementary Table 1.

### 3.4 Implementation of FL-QSAR as a prototype for collective QSAR modeling

In this study, we proposed the FL-QSAR prototype under a HFL framework for collective QSAR modeling. FL-QSAR is designed as an user-friendly prototype for predicting QSAR under HFL with pyTorch implementation. Users just need to specify the training data, test data and the client number, and then the model training, prediction and the final performance reports for each individual client as well as their collaborated one via HFL will be showed in the results files. FL-QSAR aims to help users with a better understanding of the workflow and the underline mechanism of applying HFL for QSAR modeling. FL-QSAR can be easily extended to apply other ML models for solving various drug-related learning tasks. Taking together, FL-QSAR is expected to provide great promotion for the development of collaborative drug discovery and privacy-related computing in pharmaceutical community.

## 4 Discussion

Several frameworks are presented for FL, such as FATE (https://github.com/FederatedAI/FATE), PySyft (Ryffel, et al., 2018), PaddleFL (https://github.com/PaddlePaddle/PaddleFL), etc. To the best of our knowledge, FATE is the only industrial-grade framework while other frameworks are more of theoretical value. Recently, OpenMined, which belongs to PySyft’s development community, and the PyTorch partner, have announced a plan to develop a combined platform by integrating PyTorch, PySyft and CrypTen to accelerate privacy-preserving ML. However, a FL-based collective drug discovery platform is still lacking and FL-QSAR is served as a pioneer and prototype study to inspire the related community to further investigating FL for pharmaceutical applications.

Besides HFL, VFL and FTL are also expected to have potential utilities in collaborative drug discovery. For example, different pharmaceutical companies may have their unique assay for leads discovery, therefore the distinct in-house data for the same compound in the development pipelines in different company can be generated. Integrating this information together from different institutes for the same set of compounds by VFL and FTL will accelerate the drug discovery procedure substantially, which is waiting to be explored in the future.

Another future improvement is that Federated Averaging Algorithm may be improved by aggregating parameters with a client-oriented way. Weighted average of parameters based on the training performance contributed by individual client may be an improvement direction of the Federated Averaging Algorithm, and this can be regarded as a fair way to evaluate the contribution of individual client. HFL may also be attacked by a malicious participant training a GAN (Hitaj, et al.). A malicious participant has the ability to reconstruct the training data of a given label. Therefore, how to guide the participants to make their own contributions properly and evaluate or reward their contributions fairly is important while challenging, waiting to be further explored.

In summary, biomedical community is expected to be beneficial from FL. Bio-medical data such as disease symptoms, gene sequences are very sensitive, private, and difficult to collect. If bio-medical data are collected together, the performance of ML models trained on the large scale bio-medical dataset are expected to be significantly improved, however, the collaborative and privacy-preserving learning framework applied here are needed to be carefully designed and investigated to address the facing challenges.

## Acknowledgements

The authors are grateful to the anonymous reviewers for their valuable suggestions and comments, which will lead to the improvement of this paper.

## Author contribution

Q.L. and Q.Y. conceived the study, S.Q.C., D.Y.S., G.H.C. implemented FL-QSAR. Q.L., S.Q.C. and Q.Y. wrote the manuscript with assistance from other authors.

